# Deep Learning-based Automated Rare Sperm Identification from Testes Biopsies

**DOI:** 10.1101/2021.11.14.468543

**Authors:** Ryan Lee, Luke Witherspoon, Meghan Robinson, Jeong Hyun Lee, Simon P. Duffy, Ryan Flannigan, Hongshen Ma

**Affiliations:** Department of Mechanical Engineering, University of British Columbia, Vancouver, BC, Canada; Centre for Blood Research, University of British Columbia, Vancouver, BC, Canada; Department of Urologic Sciences, University of British Columbia, Vancouver, BC, Canada; Vancouver Prostate Centre, Vancouver General Hospital, Vancouver, BC, Canada; Department of Urology, The Ottawa Hospital, Ottawa, ON, Canada; Department of Urology, Weill Cornell Medicine, New York, NY, USA; School of Biomedical Engineering, University of British Columbia, Vancouver, BC, Canada

**Keywords:** Artificial Intelligence, infertility, sperm, microTESE, non-obstructive azoospermia

## Abstract

Non-obstructive azoospermia (NOA), the most severe form of male infertility, is currently treated using microsurgical sperm extraction (microTESE) to retrieve sperm cells for *in vitro* fertilization via intracytoplasmic sperm injection (IVF-ICSI). The success rate of this procedure for NOA patients is currently limited by the ability of andrologists to identify a few rare sperm cells among millions of background testis cells. To improve this success rate, we developed a convolution neural network (CNN) to detect rare sperm from low-resolution microscopy images of microTESE samples. Our CNN uses the U-Net architecture to perform pixel-based classification on image patches from brightfield microscopy, which is followed by morphological analysis to detect individual sperm instances. This CNN is trained using microscopy images of fluorescently labeled sperm, which is fixed to eliminate their motility, and doped into testis biopsies obtained from NOA patients. We initially tested this algorithm using purified sperm samples at different imaging magnifications in order to determine the upper bounds of performance. We then tested this algorithm by doping rare sperm cells into testis biopsy samples from NOA patients and found a sperm detection F1 score of 85.2%. These results demonstrate the potential to use automated microscopy to dramatically increase the amount of testis biopsy tissue that could be comprehensively examined, which greatly increases the chance of finding rare viable sperm, and thereby increases the success rates of IVF-ICSI for couples with NOA.

## INTRODUCTION

Over 30 million men worldwide are infertile^1^. Approximately 15% of infertile men suffer from azoospermia [1], the most severe form of male infertility where there is no detectable sperm in the ejaculate. For azoospermic men, 40% have non-obstructive azoospermia (NOA) that results from a defect in sperm production. To help these couples conceive via in vitro fertilization (IVF), sperm is directly extracted from the testis through a process known as microsurgical testicular sperm extraction (microTESE). Retrieved sperm is then injected into oocytes via intracytoplasmic sperm injection (ICSI) with the aim of achieving fertilization and pregnancy that ultimately result in a healthy live birth. For NOA patients, the success rate of microTESE is ∼45%, which translates to a live birth rate of only 13%, leaving the majority of these couples with no options for fertility [2].

The primary reasons for microTESE failures include the lack of sperm production and the inability to find extremely rare sperm among the millions of testicular cells in the mTESE specimen. Biopsied tissues are mechanically and enzymatically digested into a single cell suspension. An andrologist then evaluates the suspension using a microscopy to identify and isolate ∼10 sperm from an estimated 10-50 million testicular cells – a 0.0001-0.00002% event rate. Currently, andrologists must spend hours manually searching through tens of thousands of microscopy fields in order to identify and retrieve sperm cells for NOA patients. As a result, most IVF facilities are only able to search through a fraction of the available sample, which due to extremely low event rates, have a significant risk of missing rare sperm [3]. Thus, the success rate of IVF for NOA patients are currently limited by human cognition and patience.

There have been previous attempts to improve microTESE identification of sperm with fluorescent activated cell sorting (FACS), which identified sperm in three of seven negative microTESE samples, supporting the rational for this study [3]. However, this approach requires fluorescence staining of sperm in the microTESE sample, which is incompatible with ICSI. Attempts have been made to automate sperm identification via microscopy using traditional image analysis, as well as machine learning. Traditional image analysis uses techniques such as thresholding, edge detection, morphological transformations, elliptical masks, and colour spaces to segment sperm from other cells and debris [4–9]. These approaches are typically not robust across different specimens. Some success has been found in reviewing microscopic video footage to identify sperm based on their motility [10,11]. However, sperm in testis biopsies are typically less motile than sperm in semen. Recently, deep learning has been used to directly detect sperm in semen from microscopy images, identify DNA fragmentation, and to select high-quality sperm for ICSI [12–18]. These studies demonstrate that deep learning could extract useful information from bright field images of sperm. However, the sample needs to be imaged at high magnification (>40X), which is incompatible with the need to search through all available biopsy tissue to detect rare sperm.

In order to improve the success rate of IVF-ICSI for NOA patients, we developed an artificial intelligence (AI) algorithm to automate the identification of rare sperm from semen and testis biopsy. This capability dramatically increases the amount of testis biopsy tissue that could be comprehensively examined and therefore increases the chance of finding viable sperm for ICSI. This capability would also free andrologists from an arduous task in order to make the overall ICSI process more efficient and more successful, which ultimately, will help couples devastated by NOA to achieve their dream of starting a family.

## RESULTS

### Approach

Our overall approach to automated rare sperm identification is to use a convolutional neural network (CNN) to perform pixel-based classification of microscopy images from testes biopsies (**Fig. 1**). To train the CNN, purified sperm cells are first isolated from semen samples using a swim-up procedure. The purified sperm cells are then fixed to make them non-motile and fluorescently stained using an amine-reactive dye. These sperm cells are doped into testis biopsy samples from NOA patients confirmed to have no sperm by a clinical andrologist. The doped samples are imaged using bright field and fluorescence microscopy. The fluorescence images are processed to generate a ground truth of the locations of doped sperm cells to train the CNN to perform pixel-based classification on the bright field images. The output of the CNN is a probability map of potential sperm objects. Finally, the probability distribution is processed to identify and locate individual sperm cells. The performance of our CNN model can be evaluated by comparing detected sperm from bright field images against ground truth labels obtained from fluorescence images. Prior to applying this approach to sperm doped into testis biopsy samples, we initially used this image analysis pipeline to analyze purified sperm samples without background cells in order to determine the upper bound on performance and optimal imaging magnification.

**Fig. 1.**
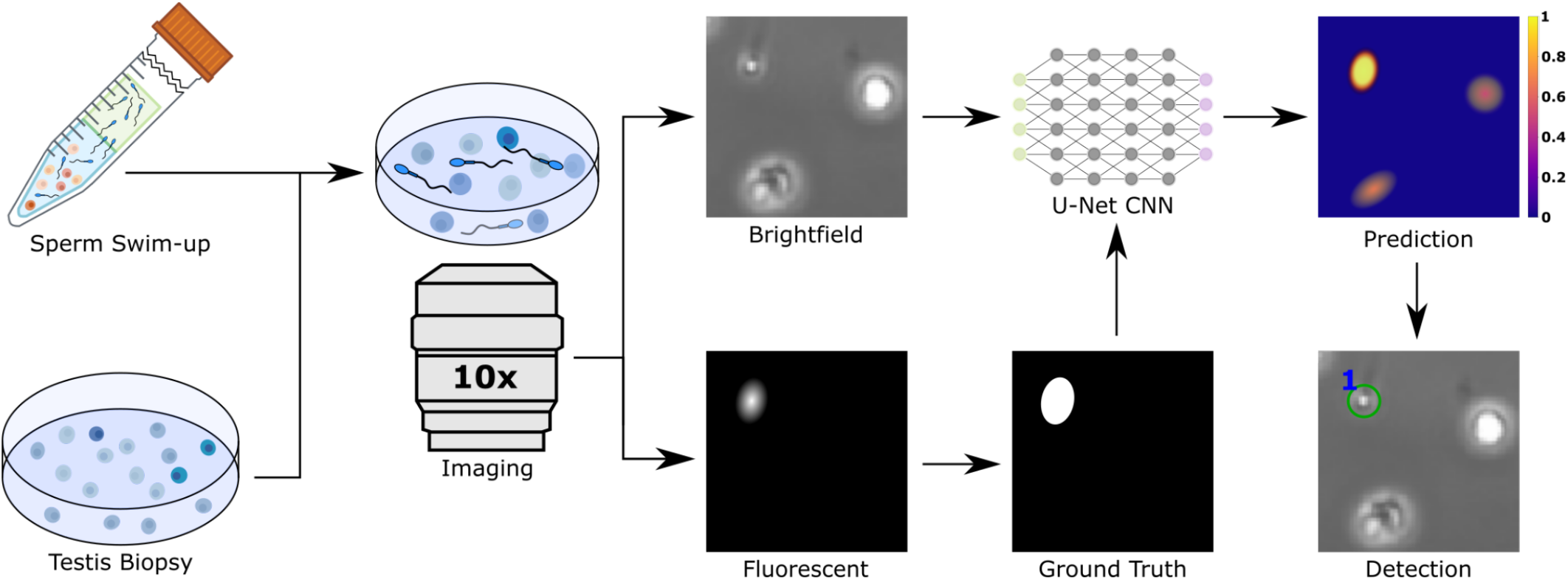
Overall approach for AI-based rare sperm detection system. Sperm purified by swim-up is fixed, fluorescently-stained, and doped into a testis biopsy, which is imaged to create a training set for deep learning. A U-Net CNN is trained using bright field images against ground truth obtained by processing the fluorescence images. The trained CNN performs pixel-based classification to obtain a probability distribution, which is analyzed to detect individual sperm.

### Data preprocessing

Bright field and fluorescence images of sperm and testis cells are acquired as 8-bit 2424×2424 pixel images from the microscope camera. We developed a Python program to resize and slice the images into 81 256×256 pixel bitmap image patches in order to eliminate low-quality regions and reduce the memory requirements for the training environment. The patches are filtered and inspected to remove well edges, as well as areas with unusually high fluorescence, large clumps, or no sperm in order to produce a high-quality training set with minimal outliers. To create the ground truth, the fluorescent images are thresholded to produce a binary image (**Fig. 2A-B**). The ground truth threshold was determined by analyzing fluorescence images of purified sperm samples to detect the sperm head. The dataset is then split into a 70%, 15%, and 15% ratio with 35,761, 7,663, and 7,663 patches for training, validation, and testing data sets respectively by random selection. This dataset split allows sufficient diversity in training images while isolating sufficient validation and testing images to produce a robust model evaluation. During model training, EarlyStopping and Checkpoint functions in Karas are applied to save and stop the training before overfitting. As image patches are fed in batches to train the model, patches are adjusted in real time using ImageDataGenerator in Karas to artificially increase the diversity of the training dataset to improve model generalizability.

**Fig. 2.**
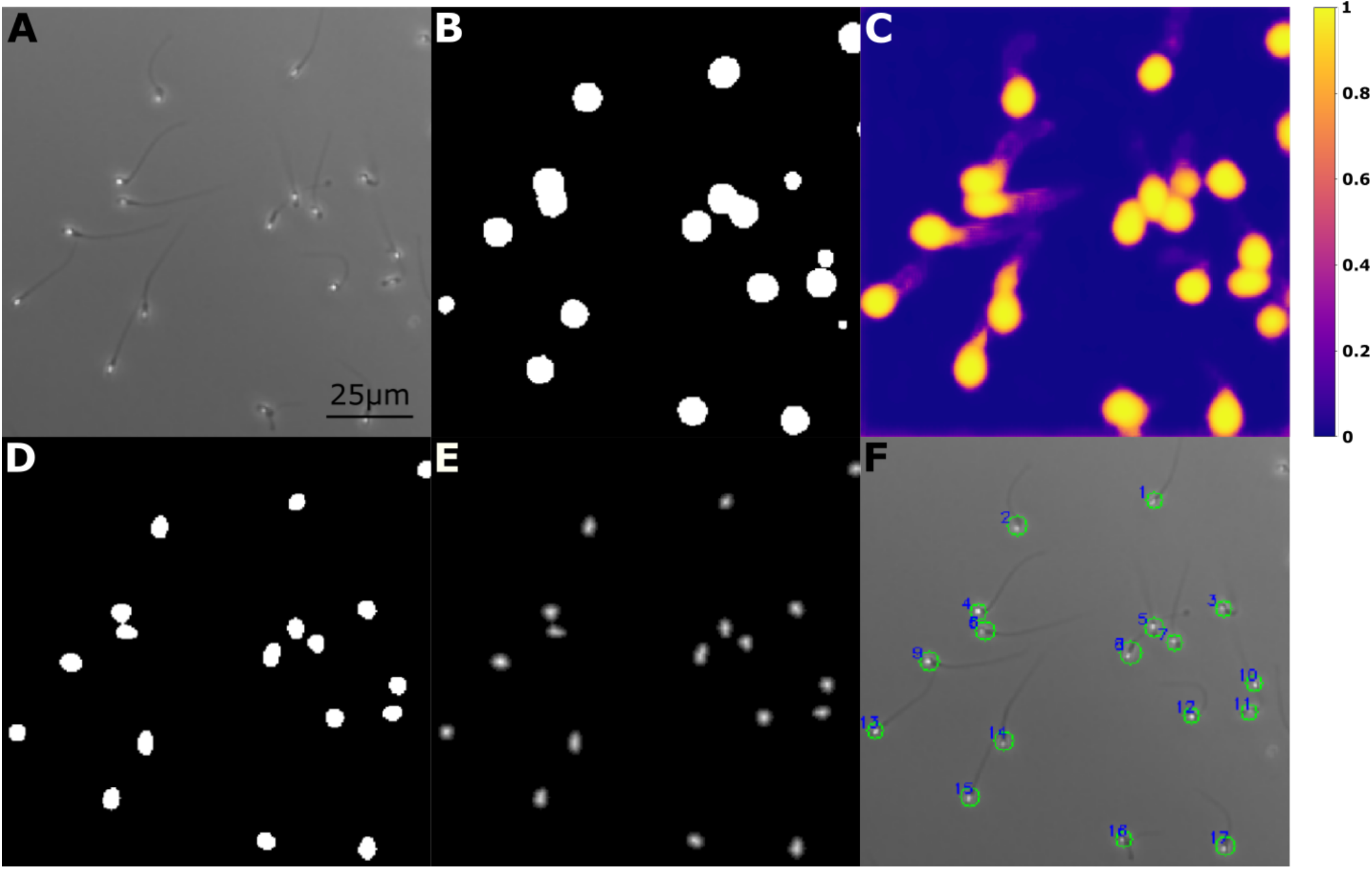
Example training images of purified sperm. **A**. An example 256×256 bright field microscopy image patch. **B**. Thresholded fluorescent image providing the ground truth label used to train the CNN model. **C**. Normalized probability distribution from the CNN. **D**. Binary mask generated by thresholding the CNN model prediction. **E**. Isolate unique sperm instances by using a watershed algorithm to determine the Euclidian distance from the background pixels. **F**. Sperm cells are located and visualized using the smallest fitted circle along with a number label for each instance.

### Network Design

We designed a CNN based on the U-Net architecture to segment images into sperm and non-sperm pixels using the Keras library in Tensorflow. U-Net is a fully convolutional encoder-decoder network architecture with a proven ability to segment medical images and conduct transfer learning with minimal data [19]. This network uses an equal number of upsampling and downsampling layers that form a symmetric contracting and expanding path. Unlike patch-based CNNs, the network can utilize the full context of each image via skip connections that concatenate features from opposing convolution and deconvolution layers **(Fig. 3)**. Several differences can be highlighted between the original U-Net architecture and our design. Specifically, we used ELU activation instead of ReLU to avoid the dying ReLU problem and dropout layers were used to regularize the model. While U-Net didn’t use dropout layers, to regularize the network, we used two different styles of dropout that were trained and compared: a constant 50% dropout, as well as increasing dropouts in the encoder and decreasing dropouts in the decoder. The model input is a three-stacked grayscale 8-bit 256×256 pixel image, such that all 2×2 max-pooling operations are applied to a layer with even height and width as suggested by U-Net, normalized about 0 between –1 and 1. We used padded 3×3 convolutions in place of padded 3×3 convolutions to maintain the input and output’s height and width, but kept the stride of 2. For weight initialization, instead of a regular normal distribution used by the U-Net, we used a truncated normal distribution to avoid rare outlier weights, but kept the standard deviation of 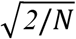, where N is the number of input units in the weight tensor. The energy function is computed pixel-wise using the two-class sigmoid function rather than the multiclass softmax function. The network uses a weighted binary cross-entropy loss function instead of an unweighted version, as well as the Adam optimizer with an initial learning rate of 0.01. This loss function increases the log loss of incorrectly segmenting sperm by the ratio of background to sperm pixels in the entire training dataset. The output segmentation is binarized into a single-layered 8-bit 256×256 pixel image for post-processing.

**Fig. 3.**
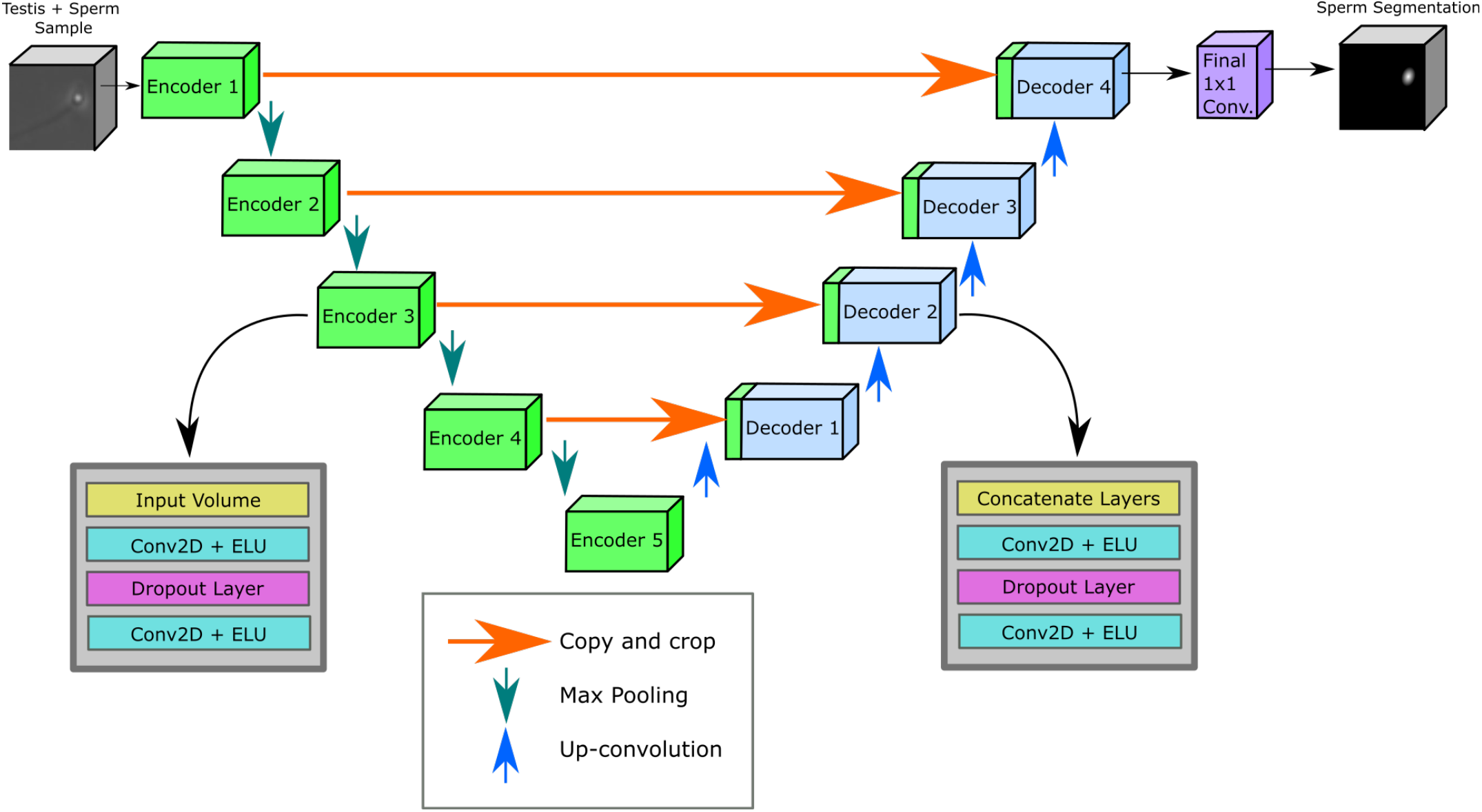
U-Net CNN. The U-Net CNN consists of a series of encoders and decoders that make use of skip connections to concatenate encoder outputs into the decoder inputs. Using dropout for regularization, max pooling to link encoders, up-convolutions to link decoders, and a 1×1 convolution with sigmoid activation to predict the pixel-based probabilities of sperm cells. Both encoders and decoders use 2D convolutions to downscale and then upscale the inputs.

### Detection

To detect each sperm and acquire their location, we post-process the CNN probability map (**Fig. 2C**) to perform instance segmentation. The CNN probability map is initially thresholded into a binary mask containing only high-confidence sperm pixels (**Fig. 2D**). The binary mask is then transformed into a Euclidean distance map using the SciPy library such that all sperm pixels are a minimum Euclidean distance from the background labels (**Fig. 2E**). The local peaks within the distance map are found using the SciKit library, with each peak identified via a unique label. The watershed algorithm is then applied to the combination of unique labels and inverse Euclidean distance map to calculate the contours surrounding each unique peak by flooding the inverse peaks to determine the contour boundaries. Each sperm instance within an image patch can then be visualized with a circle and number using the unique contours that surround each peak found using the watershed algorithm (**Fig. 2F**) to evaluate model performance. Interestingly, while sperm tails were not labeled on the ground truth images, tails are labeled on the probability maps produced by the CNN (**Fig. 2C**), which confirms that the presence of the tail is being used by the CNN to identify sperm cells.

### Testing

We measured the performance of the model by comparing the predicted sperm locations with the ground truth. If the predicted sperm location is within the approximate radius of the sperm head (six pixels for images taken using a 10× objective), the predicted sperm is labeled as a true positive. Otherwise, the predicted sperm is labeled as a false positive. All missed ground truth sperm are labeled as false negatives and no predicted sperm is counted twice. By measuring the number of true positives, false positives, and false negatives **(Fig. 4)**, we calculate the precision, recall, and F1-scores for each tested dataset to evaluate the performance of the model. The parameters of the watershed algorithm, including the binarization threshold, the maximum distance threshold to be considered a “nearby sperm”, and the confidence threshold for binarization were optimized through empirical testing with a resolution of one pixel.

**Fig. 4.**
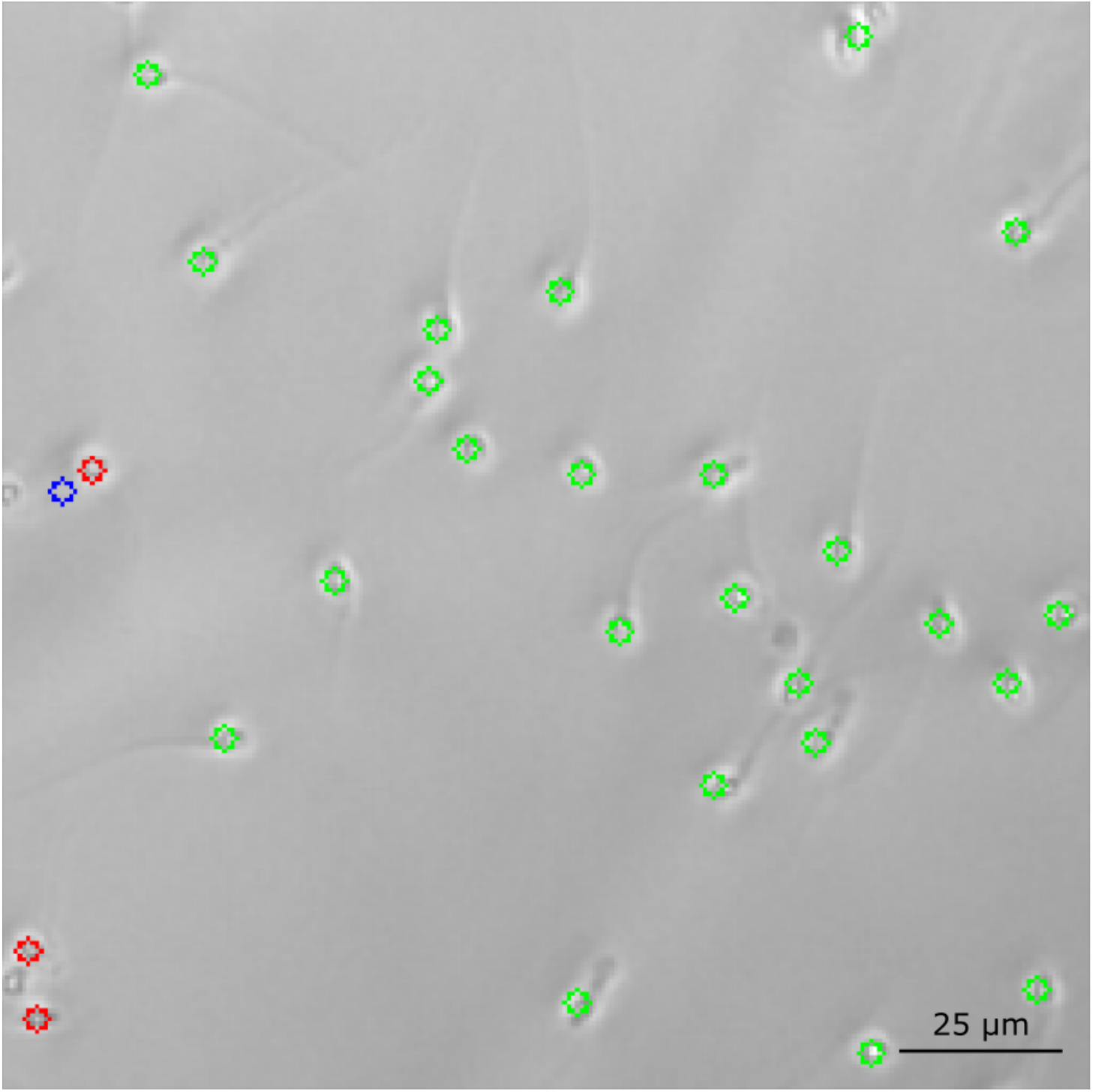
Labeled Bright Field Patch. Brightfield patch labeled with green for true positives, blue for false positives, and red for false negatives.

### Imaging Resolution

Imaging magnification has a tremendous impact on the amount of tissue that can be analyzed because the time required for microscopy increases as the inverse square of the magnification. For example, a 4×magnification objective can image 25 times the amount of biopsy tissue compared to a 20× objective. To determine the optimal imaging resolution for large area scans, we evaluated sperm detection accuracy by training our CNN using microscopy images acquired at 20×, 10×, and 4× magnification. To ensure identical images are used for this test, the 10× and 4× images are down-sampled from 20× images. At 4× magnification, the sperm head is detected using ∼8 pixels and is barely recognizable by human cognition (**Fig. 5A**). At 10× and 20×, the sperm head is detected using ∼50 and ∼200 pixels respectively, and is easily recognizable by human cognition (**Fig. 5B-C**). Training models at all three magnifications using a sample containing 68,041 sperm cells, we found precision, recall, and F1-score to be 77.8%, 69.7%, and 73.5% for 4×; 91%, 95.8%, and 93.3% for 10×; and 97.6%, 94.2%, and 95.9% for 20× magnification (**Fig. 5D**). Based on these results, we selected 10× magnification for subsequent study on testis biopsy samples because it provides reasonable imaging speed without suffering a significant loss in accuracy.

**Fig. 5.**
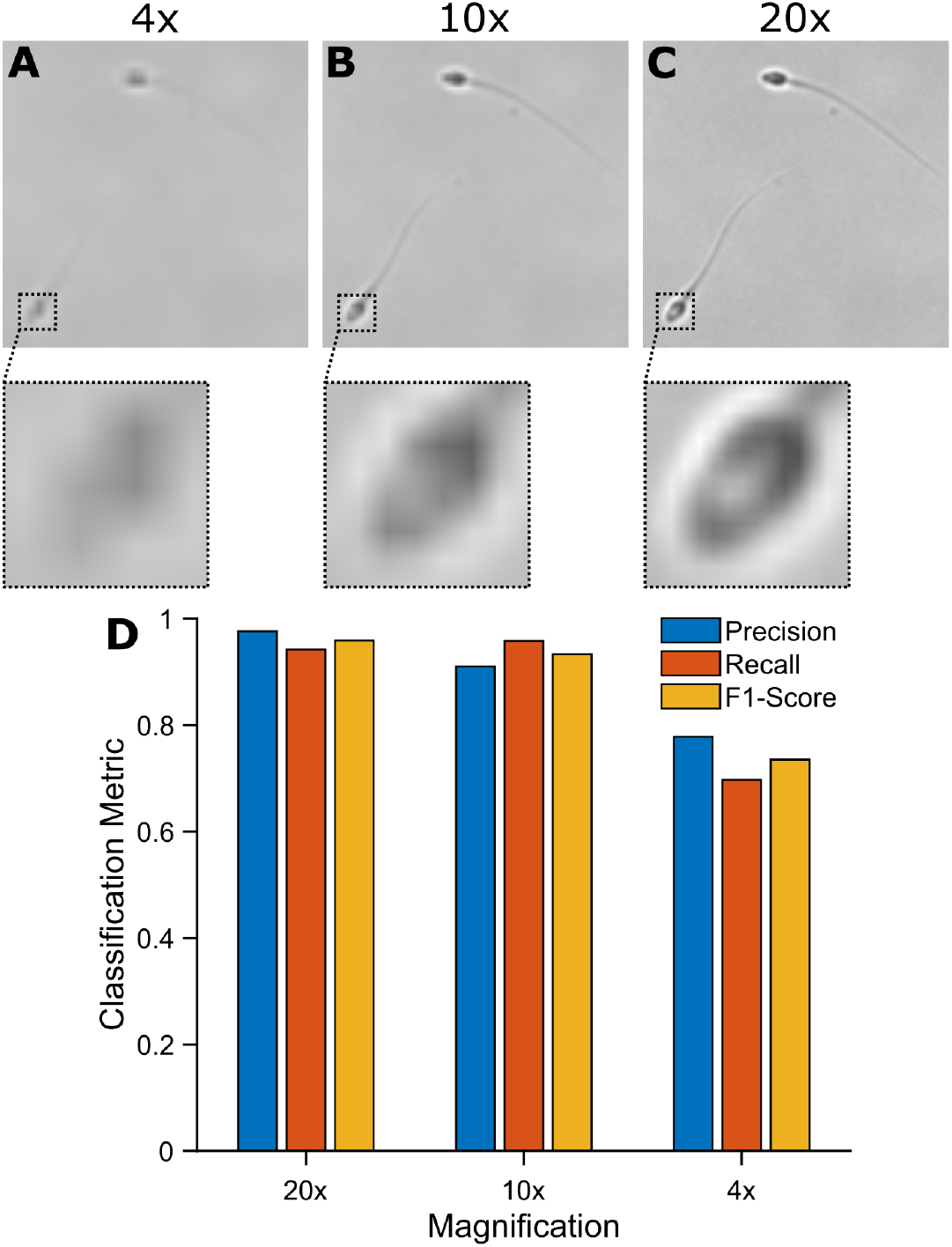
Magnification Comparison. **A**. Precision, recall, and F1-score of sperm cells detected using 4×. 10×, and 20× objectives using sperm-only data. **B-D**. Images acquired using 4×, 10×, and 20× objectives including magnified regions around a single sperm head.

### Sperm Identification in Testis Biopsy

We tested our approach for identifying sperm in testis biopsy imaged using the 10× microscope objective. We trained a new CNN model by doping purified sperm cells doped into testis biopsy from NOA patients, who have been established to be azoospermic. As before, the purified sperm is stained using an amine reactive dye to create a ground truth **(Fig. 6A & 6B)**. The testis biopsy sample is then imaged in both bright field (**Fig. 7A**) and fluorescence. After preprocessing to eliminate low-quality images, the fluorescence images are binarized using an empirically tested threshold dependent on microscopy gain, exposure time, and microscope, to provide ground truth images containing the sperm head for training the CNN model to detect sperm in the bright field images (**Fig. 7B**). The output of the CNN is a probability map of the likelihood of pixels containing sperm (**Fig. 7C**). To obtain the Euclidean distance map using the distance_transform_edt function in SciPy, the probability map must be binarized into an input binary image of positive and background pixels using a confidence threshold optimized through testing (**Fig. 7D**). After transforming the binary patch into a Euclidean distance map (**Fig. 7E**), local maxima points are identified using the peak_local_max function in SciPy, and then uniquely labeled using the label function in SciPy. A watershed algorithm is used to determine the contour that belongs to each detected sperm instance by filling the area of the binarized prediction (**Fig. 7D**) beginning at the sperm and stopping when it meets the boundary of another sperm instance. As before, the sperm tails are detected in the probability map even though the training images had no sperm tails (**Fig. 7C & 7F**).

**Fig. 6.**
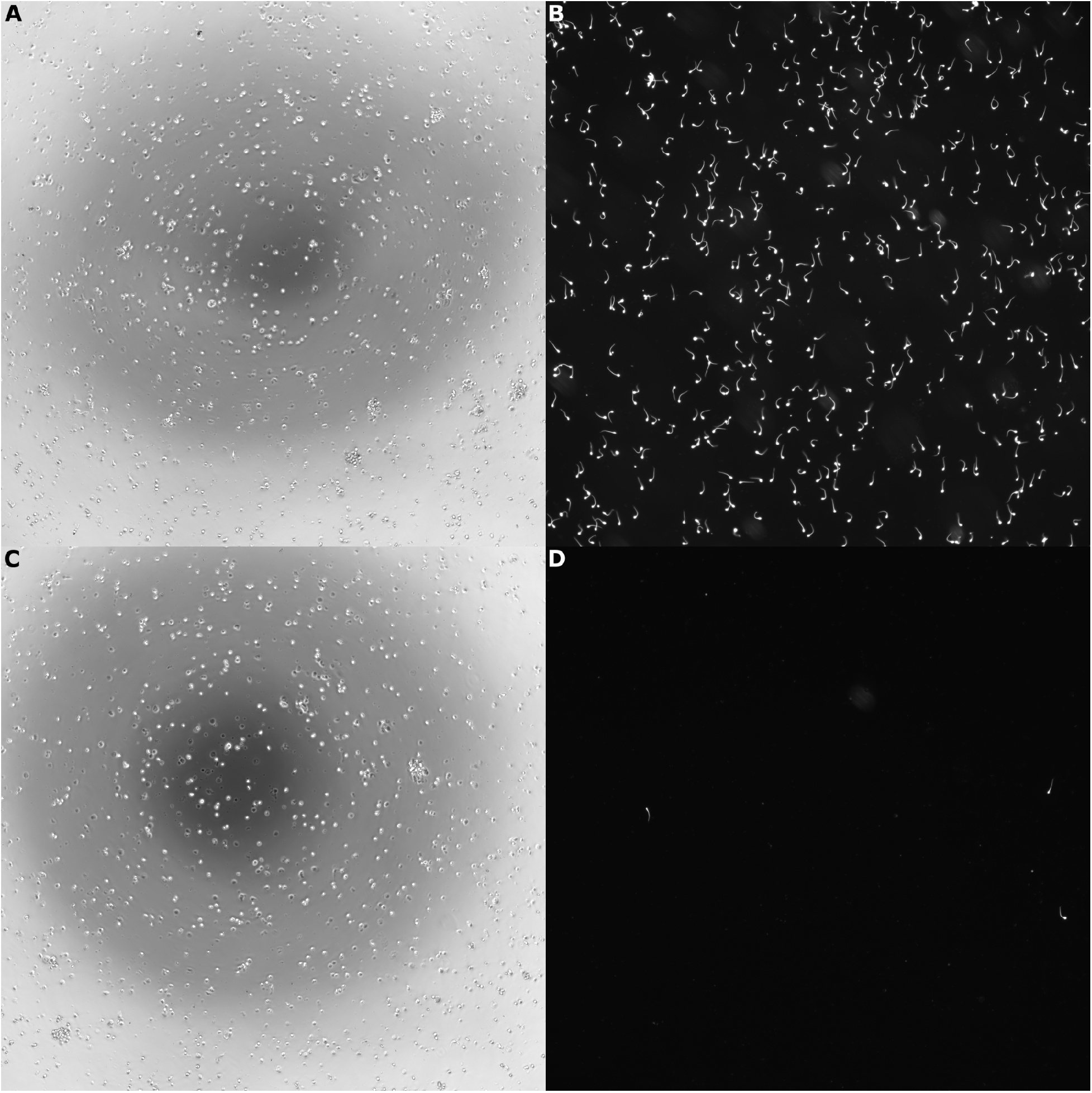
Example bright field and Fluorescent Images of sperm. **A**. An example 2024×2024 pixel bright field image acquired at 10×. **B**. An example 2024×2024 pixel fluorescent image acquired at 10×. **A**. An example 2024×2024 pixel bright field image of rare sperm acquired at 10×. **B**. An example 2024×2024 pixel fluorescent image of rare sperm acquired at 10×.

**Fig. 7.**
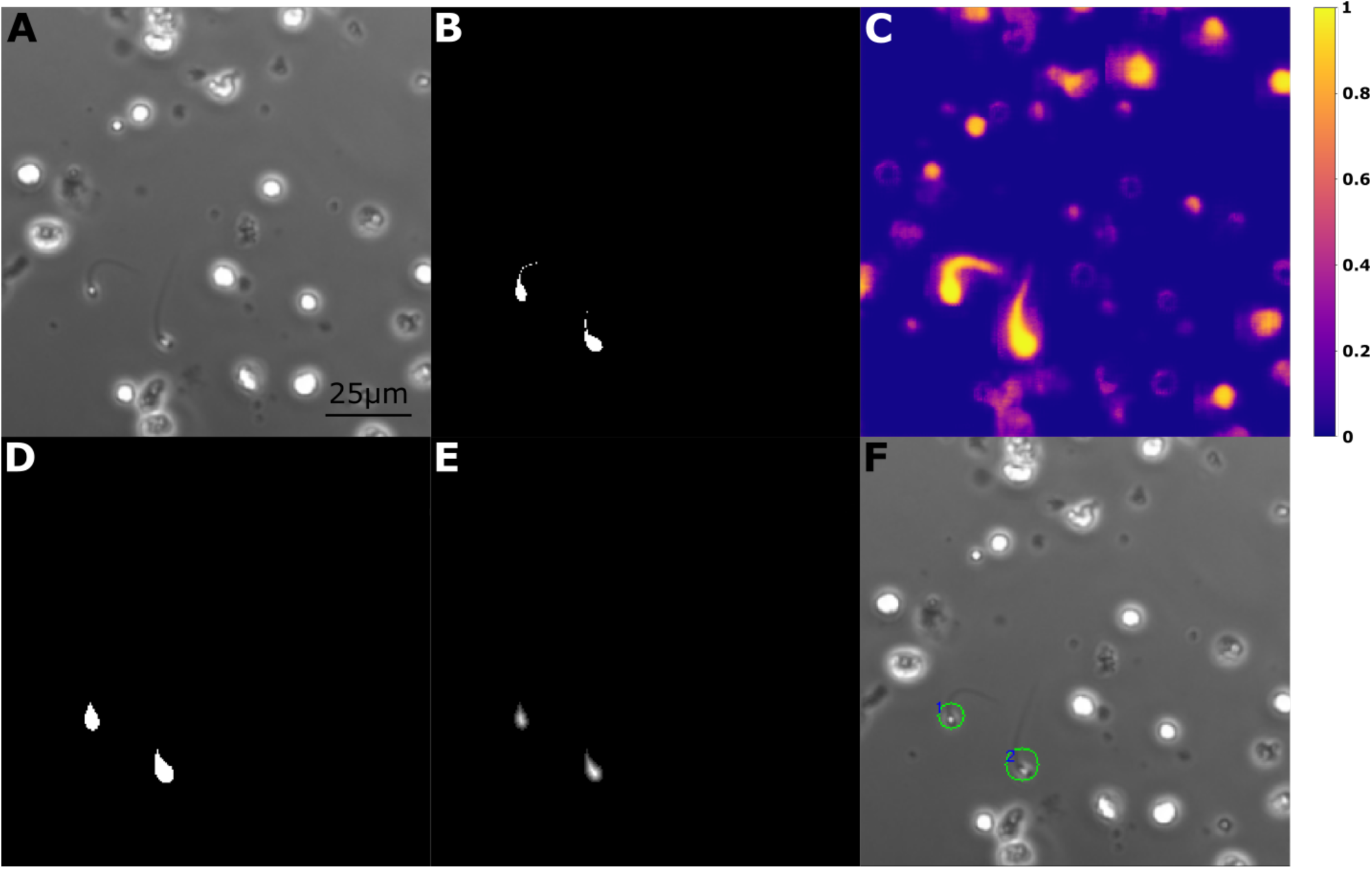
Example training images sperm doped into testis biopsy. **A**. An example 256×256 bright field microscopy image patch. **B**. Thresholded fluorescent image providing the ground truth label used to train the CNN model. **C**. Normalized probability distribution provided by the CNN model. **D**. Binary mask generated by thresholding the CNN model prediction. **E**. Isolate unique sperm instances using a watershed algorithm to determine the Euclidian distance from the background pixels. **F**. Sperm are located and visualized using the smallest fitted circle along with a number label for each instance.

To optimize sperm detection from the probability map (**Fig. 7C**), we vary the binarization threshold and the resolution of the watershed algorithm’s output contour radius with the goal of obtaining the greatest F1-score. The binarization threshold controls the required model prediction confidence reflected in its output probability maps for cells to be possibly a sperm. The minimum contour radius directly controls the confidence area for a cell to be labeled as a sperm. To optimize these two parameters, we consider the task of identifying rare sperm for ICSI, which requires a balance of precision and recall, therefore F1-score is the most relevant metric. Using detected sperm locations, we measure the model performance on the validation dataset, which is then used to optimize the aforementioned binarizing threshold with a resolution of 0.01 and minimum contour radius with a resolution of 1 pixel. The combination of parameters that yielded the highest F1-score was selected to be the optimal model. We optimized F1-scores for two dropout styles: increasing/decreasing dropouts and a constant 50% dropout. The former achieved 78.7% (**Fig. 8**) and the latter 74.8%. In summary, our CNN can successfully identify sperm in testis biopsy samples with an F1-score of 78.7%.

**Fig. 8.**
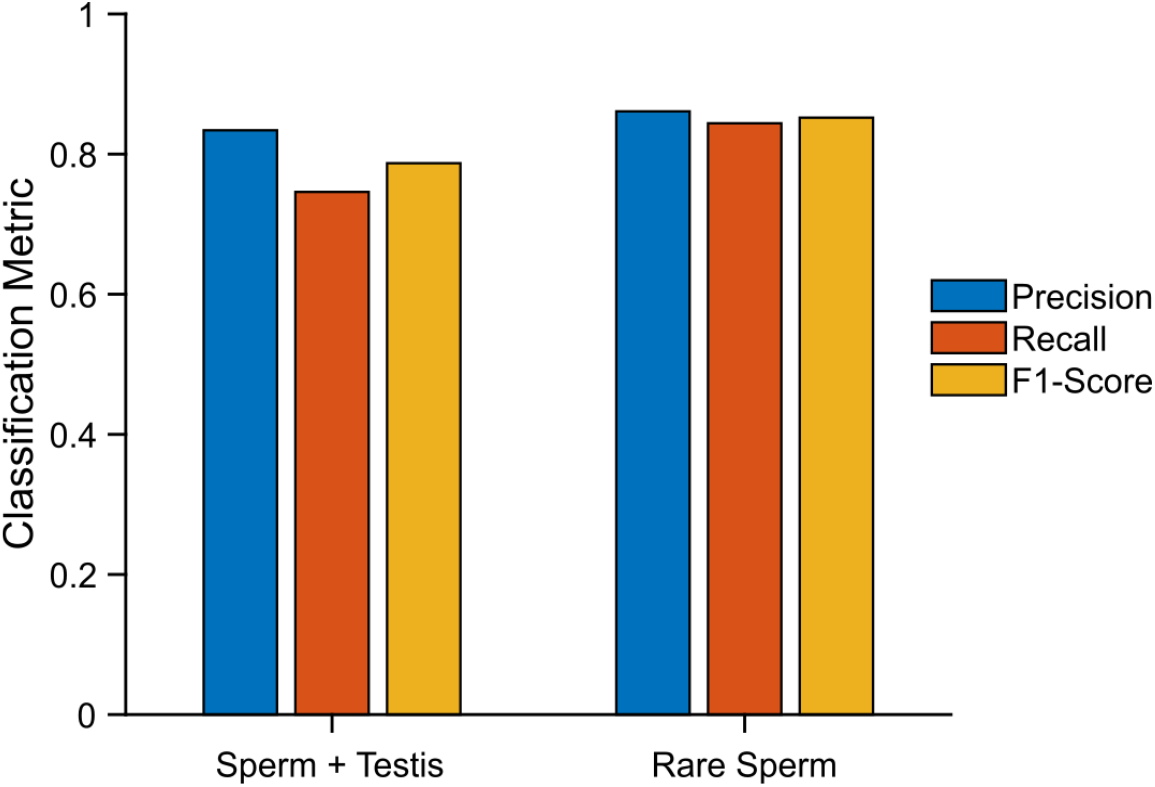
Final metrics. Precision, recall, and F1-scores for detecting sperm in testis biopsy tissue for both abundant sperm and rare sperm images using the same trained model.

Additionally, to establish the consistency of our results, we validated our results using five-fold cross-validation. The training set was split into five equally sized and randomly distributed groups of images. We trained the U-Net using four of the groups and tested it on the fifth group. All five groups have been tested after repeating this process four times. The final metrics had a precision, recall, and F1-score of 84 ± 2.28%, 72.7 ± 2.41%, and 77.9 ± 0.72% respectively, which is similar to the performance on the initial test set.

### Rare Sperm Identification in Testis Biopsy

To investigate the use case of identifying rare sperm in testis biopsies from NOA patients where there could be only a few sperm cells among millions of testis cells, we simulated this clinical application using sperm doped artificially into testis biopsy samples that were inspected by andrologists to contain no sperm. We first purified donor sperm from semen samples using a swim-up assay. The purified sperm cells are fixed, permeabilized, and stained using an amine-reactive dye. The stained sperm are then doped in low numbers into disassociated testis biopsy cells in microwell plates. Each well had 10 to 200 sperm cells and approximately 30,000 to 50,000 cells from the testis biopsy **(Fig. 6C & 6D)**. The model was previously trained by doping 3,000 to 6,000 sperm per well, and was not retrained before testing on the rare sperm dataset. Our model was able to detect 2,969 sperm cells out of a total of 3,517 sperm in the 7,985 image patches imaged from 38 wells. This result slightly outperformed the expected 2,624 sperm cells based on the 74.6% recall from sperm identification in testis biopsy. The overall model performance had a recall of 84.4%, precision of 86.1%, and an F1-score of 85.2% (**Fig. 8**), showing that a low frequency of sperm does not negatively affect the sperm detection in the model.

## DISCUSSION AND CONCLUSIONS

Identifying rare sperm from testis biopsy of NOA patients for ICSI is an arduous task that is currently limited by human cognition and patience. Automating this task using computational image analysis provides an opportunity to dramatically increase the amount of tissue that can be assessed, and thereby increasing the chance of finding rare sperm that could be used for ICSI. In this study, we developed a CNN based on the U-Net architecture to identify sperm via pixel classification of microscopy images of testis biopsies. We trained the CNN using purified donor sperm that are fluorescently stained and doped into testis biopsy samples from NOA patients. The donor sperm was fixed using formalin to ensure that sperm are non-motile for imaging, which interestingly, did not impede their detection by the CNN. Using the trained CNN to analyze bright field microscopy images of testis biopsy tissue produced a probability map of potential sperm, which is then analyzed to identify specific instances of sperm.

We initially trained the CNN using microscopy images of purified sperm without testis cells in order to determine the upper bound on classification accuracy. We also used this data set to determine the optimal imaging magnification since computational analysis of microscopy images could be performed at a lower magnification than the 40× microscope objective used by human reviewers. Reducing the objective magnification greatly increases the amount of biopsy tissue that could be examined because the imaging area increases as the inverse square of the magnification. We found that the CNN is capable of identifying sperm at 4× magnification where each sperm head is imaged using only ∼8 pixels, with a classification F1-score of 73.5%. However, to ensure robustness, we decided to image using a 10× objective, which provided a classification F1-score of 93.3% with reasonable throughput. Training and testing our CNN on sperm doped into testis biopsy samples at 10x magnification, we found a classification F1-score of 78.7% for samples doped with abundant sperm and 85.2% for samples doped with rare sperm. The greater F1-score for rare sperm samples compared to abundant sperm samples suggest that the sperm detection in abundant sperm samples may be impaired by high sperm density.

The ability to perform automated detection of rare sperm from brightfield microscopy images is a tremendous asset for treating male infertility in NOA patients because identifying viable sperm is a key limiting factor for initiating IVF-ICSI. Therefore, increasing the amount of tissue that can be evaluated using microscopy greatly increases the potential of finding rare sperm cells, which will improve the success rates of IVF-ICSI for NOA patients. Automating sperm detection will also free up andrologists from an arduous task to reduce the total human hours that must be dedicated for each IVF-ICSI procedure.

The key novelty of this work is the approach for training a CNN for rare cell identification by creating training data using mixtures of target and non-target cells, and then using the U-Net architecture to generate a probability map of target cells. Whereas previous realizations of U-Net for segment typically measured performance using intersection-over-union between prediction and ground truth pixels, performance in this instance is better measured using object matching between prediction and ground truth. We believe this approach can be applied broadly for identifying rare cells in microscopy images of tissues.

## METHODS

### Sample Preparation

The study used fresh donated semen from normal donors, as well as mTESE samples from biobanked specimens at our institution. Within 30 minutes of a fresh semen donation, the sample was transferred to a 15 mL centrifuge tube and a needle was used to pop all bubbles on the sample. The tube is then secured at 45° and 2 mL of the swim-up medium, made from 38 mL DMEM/F12 (11320-033, Gibco), 100 µL Lactate (L4263, Sigma), 10 mL of 50 mg/mL stock has (A1653, Sigma), 0.5 mL HEPES buffer solution (15630-080, Gibco), 272 µL Sodium pyruvate (S8636, Sigma) was gently layered on top of the semen sample to perform the sperm swim-up, where the motile sperm swim upwards past the boundary into the swim-up medium, leaving behind other cells, debris, and non-motile sperm in the semen. For washes, all centrifugation was spun at 400 g for 5 min. After 20 min of the swim-up process, the top 70% of the uniform unclouded swim-up medium was slowly extracted using a large-bore pipette tip to avoid suction of the semen sample. To confirm a successful swim-up, 10 µL of the swim-up medium was viewed under a microscope on a glass slide to ensure it only contains sperm cells with negligible debris or non-sperm cells. Sperm cells are washed and resuspended in 500 µL of PBS and then 10 µL/mL formalin is added to fix the sperm. 15 min after adding the formalin, the cells were washed and resuspended in 500 µL of 0.1% Triton X-100 for permeabilization. 30 min after resuspension in Triton X-100, the sample was washed again and resuspended in 500 µL of PBS. Sperm cells were stained with 1 µL/mL LIVE/DEAD™ Fixable Aqua Dead Cell Stain (L34957, Invitrogen) and left at room temperature (with container wrapped in aluminum foil) for 30 min to allow the stain to fully permeate. An amine-reactive dye is necessary to prevent dye leakage into the testis cells using to the dye’s covalent bonds. After two more sperm cell washes in PBS, both sperm cells and mTESE cells were serially diluted and aliquoted at low density into Greiner Sensoplate 96-well glass-bottom multiwell plates (M4187-16EA, Sigma-Aldrich) such that there was minimal overlap between cells under the microscope after plate centrifugation. Before imaging, each well is layered with 15 µL of Ovoil oil (10029, Vitrolife), a total volume of 120 µL.

### Microscopy

Microscopy imaging was performed using a Nikon Ti-2E inverted fluorescence microscope. Microscope objectives included a Nikon CFI Plain Fluor 4×, 10×, and 20× objectives. Image acquisition was performed using a 14-bit Nikon DS-Qi2 CMOS camera. Images were acquired in brightfield with phase contrast, as well as in fluorescence in the mCherry channel (Nikon C-FLL LFOV, 562/40 nm excitation, 641/75 nm emission, and 593 dichroic mirror). Brightfield imaging was illuminated by the built-in Ti-2E LED. Epifluorescence excitation was performed using a 130 W mercury lamp (Nikon C-HGFI). Exposure, gain, and vertical offset were automatically determined using built-in NIS functions to avoid user bias. Cells were imaged in Greiner Sensoplate 96-well glass-bottom multiwell plates (M4187-16EA, Sigma-Aldrich). Cell concentrations in wells were diluted down to ∼30,000 to ∼50,000 cells to lower overlap between cells. NIS Jobs was used to automatically capture 21 images on each of the two channels for each well. The images were exported from NIS to standard 8-bit TIFF format. Of all imaged cells, sperm cells consisted of 3 - 25% for the training set and less than 1% for the rare sperm testing set, varied for the training set for diversification of the training dataset.

### Date preprocessing

TIFF files exported from Nikon NIS were preprocessed using a custom Python script utilizing SciKit and OpenCV libraries. The script extracted and resized 2424 × 2424 pixel TIFF images to 2304 × 2304. The resized images were each sliced into 81 256 × 256 pixel image patches. Image patches were filtered to remove well edges and out-of-focus images. BF images outside of the well or on the well edges where the images are out of focus were removed from the dataset. Subsequently, some out-of-focus labels in the fluorescence channel were filtered out by removing all labels that contain more than a fifth of the pixels are sperm. Additionally, the BF patches were filtered for undesirable lighting conditions and well edges by ensuring their average pixel intensity ranged between 70 and 230, or were removed otherwise. BF images are converted to 3 channels to match the model input. This process is repeated for each stack of TIFF images.

### Convolutional Neural Network Model

The CNN based on the U-Net architecture was designed in Python using Tensorflow’s Keras library. The network accepts a 3-channel input of size 256 × 256 pixel and outputs a single channel output of size 256 × 256 pixel. All input data is normalized to between −1 to 1, inclusive. All 2D convolutions have a kernel size of 3×3, same padding, he_normal initialization, and ELU activation. The filter size begins at 16 and doubles after every 2nd convolution until the end of the encoder segment after the 10th convolution with a filter size of 256. Each pair of 2D convolutions in the encoder is followed by a 2×2 max pooling operation. The decoder starts with a series of up convolutions with a kernel size of 2×2 and 128 filter size, concatenation with the same shaped output from the encoder, and then pairs of convolutions with the same filter size, halving until the end of the encoder after every second 2D convolution. Between every pair of 2D convolutions, we tested both an increasing dropout that increments from 0.1 until 0.5 and decrements back down to 0.1 after each pair of convolutions in intervals of 0.1, as well as a constant dropout of 0.5. The output 1×1 convolution with sigmoid activation outputs confidence values between 0 and 1.

### Training Environment

The software was run on a single computer operating Windows 10 with an Intel i7-7700 running at 3.6 GHz. There was 32 GB of DDR4 RAM running at 2400 MHz. The graphics card was a 2 GB GT 730. The training was done in Python 3.7 utilizing the Keras 1.1.2 library.

### Training

The network was trained on 10 epochs with Adam optimization and a learning rate of 0.001 with a batch size of 8. The EarlyStopping and Checkpoint functions in Keras were used to save copies of the model before the 10^th^ epoch to minimize overfitting. The weighted binary cross entropy loss function was used to train the model, where the log loss from false negatives is scaled up by the ratio of background pixels and sperm pixels over the entire training dataset. Accuracy and dice loss were used to observe model performance during training. Using the ImageDataGenerator in Keras, data augmentation was applied to all training inputs, included rotations of 45 degrees in either direction, zooming in or out between 0.8× to 1.2× magnification, vertical and horizontal image shifting of up to 0.4× the width or height, brightness adjustments of 0.8× to 1.2×, vertical and horizontal flips, and a reflecting fill when augmentations leave blank space.

### Detection

The single-channel 256 × 256 pixel output from the model prediction is binarized using a threshold optimized on the validation set during testing. Using the distance_transform_edt function in SciPy, the binary image is transformed into a map where all sperm pixels are the Euclidean pixel distance away from the background pixels. All the peaks within the Euclidean map are found with a minimum 6 pixel separation distance between peaks using peak_local_max function in SciKit and each peak is uniquely labeled using the label function in SciPy before passing both the labels and an inverted version of the Euclidean distance map into the watershed algorithm. For each unique label, the watershed algorithm determines the contour surrounding each label. Each sperm instance is visualized on the original BF image using a minimum enclosing circle and number around the predicted sperm head.

## References

[1] Cullen I, Muneer A. Surgical Sperm Retrieval and MicroTESE. In: Allahbadia GN, Ata B, Lindheim SR, Woodward BJ, Bhagavath B, editors. Textbook of Assisted Reproduction, Singapore: Springer; 2020, p. 193–202. https://doi.org/10.1007/978-981-15-2377-9_23.

[2] Vloeberghs V, Verheyen G, Haentjens P, Goossens A, Polyzos NP, Tournaye H. How successful is TESE-ICSI in couples with non-obstructive azoospermia? Human Reproduction 2015;30:1790–6. https://doi.org/10.1093/humrep/dev139.

[3] Mittal S, Mielnik A, Bolyakov A, Schlegel P, Paduch D. Pd68-01 pilot study results using fluorescence activated cell sorting of spermatozoa from testis tissue: a novel method for sperm isolation after tese. Journal of Urology 2017;197:e1339–e1339. https://doi.org/10.1016/j.juro.2017.02.3129.

[4] Medina-Rodríguez R, Guzmán-Masías L, Alatrista-Salas H, Beltrán-Castañón C. Sperm Cells Segmentation in Micrographic Images Through Lambertian Reflectance Model. In: Azzopardi G, Petkov N, editors. Computer Analysis of Images and Patterns, Cham: Springer International Publishing; 2015, p. 664–74. https://doi.org/10.1007/978-3-319-23117-4_57.

[5] Park KS, Yi WJ, Paick JS. Segmentation of sperms using the strategic Hough transform. Ann Biomed Eng 1997;25:294–302. https://doi.org/10.1007/BF02648044.

[6] Hidayatullah P, Zuhdi M. Automatic sperms counting using adaptive local threshold and ellipse detection. 2014 International Conference on Information Technology Systems and Innovation (ICITSI), 2014, p. 56–61. https://doi.org/10.1109/ICITSI.2014.7048238.

[7] Chang V, Saavedra JM, Castañeda V, Sarabia L, Hitschfeld N, Härtel S. Gold-standard and improved framework for sperm head segmentation. Computer Methods and Programs in Biomedicine 2014;117:225–37. https://doi.org/10.1016/j.cmpb.2014.06.018.

[8] Nafisi V, Moradi M, Nasr-Esfahani MH. Sperm Identification Using Elliptic Model and Tail Detection 2004;6.

[9] Mahdavi HS, Monadjemi A, Vafae A. Sperm detection in video frames of semen sample using morphology and effective ellipse detection method. J Med Signals Sens 2011;1:206–13.

[10] Ilhan HO, Aydin N. Smartphone based sperm counting - an alternative way to the visual assessment technique in sperm concentration analysis. Multimed Tools Appl 2020;79:6409–35. https://doi.org/10.1007/s11042-019-08421-3.

[11] Li Q, Chen X, Zhang H, Yin L, Chen S, Wang T, et al. Automatic human spermatozoa detection in microscopic video streams based on OpenCV. 2012 5th International Conference on BioMedical Engineering and Informatics, 2012, p. 224–7. https://doi.org/10.1109/BMEI.2012.6513003.

[12] Nissen MS, Krause O, Almstrup K, Kjærulff S, Nielsen TT, Nielsen M. Convolutional Neural Networks for Segmentation and Object Detection of Human Semen. In: Sharma P, Bianchi FM, editors. Image Analysis, Cham: Springer International Publishing; 2017, p. 397–406. https://doi.org/10.1007/978-3-319-59126-1_33.

[13] Ghasemian F, Mirroshandel SA, Monji-Azad S, Azarnia M, Zahiri Z. An efficient method for automatic morphological abnormality detection from human sperm images. Computer Methods and Programs in Biomedicine 2015;122:409–20. https://doi.org/10.1016/j.cmpb.2015.08.013.

[14] Carrillo H, Villarreal J, Sotaquira M, Goelkel A, Gutierrez R. A Computer Aided Tool for the Assessment of Human Sperm Morphology. 2007 IEEE 7th International Symposium on BioInformatics and BioEngineering, 2007, p. 1152–7. https://doi.org/10.1109/BIBE.2007.4375706.

[15] Bijar A, Benavent A, Mikaeili M, Khayati R. Fully automatic identification and discrimination of sperm parts in microscopic images of stained human semen smear. Journal of Biomedical Science and Engineering 2012;05:384–95. https://doi.org/10.4236/jbise.2012.57049.

[16] Amann RP, Waberski D. Computer-assisted sperm analysis (CASA): Capabilities and potential developments. Theriogenology 2014;81:5-17.e3. https://doi.org/10.1016/j.theriogenology.2013.09.004.

[17] McCallum C, Riordon J, Wang Y, Kong T, You JB, Sanner S, et al. Deep learning-based selection of human sperm with high DNA integrity. Commun Biol 2019;2:1–10. https://doi.org/10.1038/s42003-019-0491-6.

[18] Wang Y, Riordon J, Kong T, Xu Y, Nguyen B, Zhong J, et al. Prediction of DNA Integrity from Morphological Parameters Using a Single-Sperm DNA Fragmentation Index Assay. Advanced Science 2019;6:1900712. https://doi.org/10.1002/advs.201900712.

[19] Falk T, Mai D, Bensch R, Çiçek Ö, Abdulkadir A, Marrakchi Y, et al. U-Net: deep learning for cell counting, detection, and morphometry. Nat Methods 2019;16:67–70. https://doi.org/10.1038/s41592-018-0261-2.

